# The ratio of cytotoxic lymphocytes to M2-like macrophages is prognostic in immunogenic tumors

**DOI:** 10.1101/2021.03.24.436814

**Authors:** Artur Mezheyeuski, Max Backman, Johanna Mattsson, Alfonso Martín-Bernabé, Chatarina Larsson, Ina Hrynchyk, Klara Hammarström, Simon Ström, Joakim Ekström, Siarhei Mauchanski, Salome Khelashvili, Margrét Agnarsdóttir, Per-Henrik Edqvist, Jutta Huvila, Ulrika Segersten, Per-Uno Malmström, Johan Botling, Björn Nodin, Charlotta Hedner, David Borg, Jenny Brändstedt, Hanna Sartor, Karin Leandersson, Bengt Glimelius, Anna Portyanko, Fredrik Ponten, Karin Jirström, Patrick Micke, Tobias Sjöblom

## Abstract

Immune cells in the microenvironment shape tumor development and progression. The prognostic value of T-cell-based immune scores exceeds those of clinical parameters in colon cancer, but reflects only a part of the anti-tumor immune response. Here, we assessed 15 distinct immune cell classes and identified a simple prognostic signature based on pro- and anti-tumoral immune cells in the tumor microenvironment. The ratio of cytotoxic lymphocytes to tumor supportive macrophages predicted survival better than the state-of-art immune score in colon cancer and had the highest relative contribution to survival prediction when compared to established clinical parameters. This signature was prognostic also in other cancers with high mutation burden, such as those of the lung, bladder, esophagus, and melanomas, supporting broad clinical applicability.

**One Sentence Summary:** The CD8/M2 ratio in tumor tissue defines prognosis in immunogenic cancers

## Main text

Colorectal cancer (CRC) is the fourth most common and the second most lethal type of cancer (*1*). The traditional TNM classification, cancer gene mutations and expression profiles identify more homogeneous subgroups among the intrinsically heterogeneous CRC tumors (*2*). Recently, the immune response was acknowledged as a prognostic factor in CRC (*3*). An immune scoring system that evaluates the abundance of CD3^+^ and CD8^+^ T cells in resected tumors was recently validated as an independent prognostic factor in colon cancer stage I-III, surpassing established clinical parameters such as T and N stage (*4, 5*). The Immunoscore®, although of proven validity, quantitates only a limited subset of anti-tumoral immune cells, but a growing body of evidence supports association of additional immune cell types, including B cells and NK cells, with better outcome (*6–9*). In addition, T-regulatory lymphocytes and M2-polarised macrophages residing in the tumor microenvironment have been connected to tumor progression (*10*), suggesting pro-tumoral effects and prognostic potential of these immune suppressive cells. The aims of this study were (1) to generate a comprehensive overview of the immune landscape in CRC by *in situ* analysis of 15 distinct subclasses of T- and B-lymphocytes, myeloid cells and NK cells, (2) to identify the immune cell signature with the highest prognostic value, and (3) to assess the prognostic ability of this signature in other tumor types.

To quantitate the different immune cell subsets, we performed immunohistochemistry (IHC) and multispectral imaging enabling multiplex labeling of markers in tumor tissue. We used two IHC panels, each consisting of antibodies to five immune markers, for visualization of adaptive and innate immune cells. After cell segmentation of digitized tissue sections, the co-expression patterns of these markers allowed for immune cell classification into distinct subgroups (see (*9, 11, 12*) and Supplementary Materials and Methods) (Fig 1A). The major immune cell lineages were defined by single markers (CD4, CD8, CD45RO, CD68 and CD163). By marker co-expression we identified memory CD4 and CD8 lymphocytes, classical CD4^+^ T-regulatory and CD8^+^ Treg cells. For natural killer (NK) cells, we required co-expression of two markers (CD56 and NKp46) to classify a cell as NK, along with CD3 expression to classify as NK T (NKT) cell. Finally, the monocyte/macrophage lineage was sub-divided into M1-like macrophages (CD68^+^CD163^−^), M2-like macrophages (CD68^+^CD163^+^) and CD68^−^CD163^+^ cells.

**Fig. 1.**
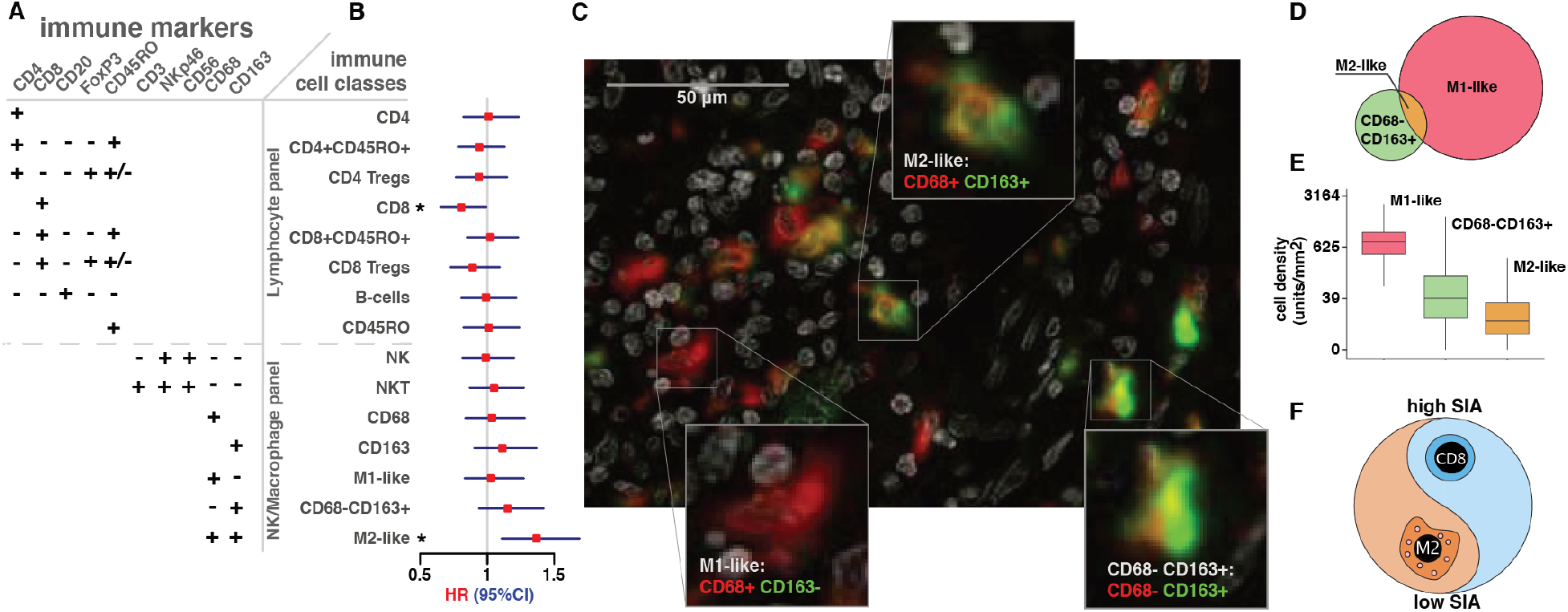
Prognostic value of CD8+ T cells and CD68+/CD163+ macrophages revealed by comprehensive characterization of immune cell subsets in 286 therapy-naïve colon cancers. Immune marker combinations in IHC panels define classes and subclasses of immune cells. Forest plot of univariate associations of tissue immune cell densities translated into three-level categorized values, with OS in patients of stage I-III. Filled squares, hazard ratios (HR); whiskers, 95% confidence intervals (CI), \**p*<0.05 (Cox regression). **(C)** Representative multiplex macrophage marker staining of colon cancer tissue. Expression of two markers, CD68 (red) and CD163 (green) with nuclear DAPI staining (white), visualized in pseudocolors, identified three cell types (insets), M1-like macrophages, M2-like macrophages and CD163^+^CD68^−^cells. **(D)** Venn diagram of the counts of cells in the entire cohort expressing CD68 only (red, *n*=9.0×10^5^), CD163 only (green, *n*=1.9×10^5^) or both markers (gold, *n*=4.4×10^4^). **(E)** Density of three macrophage subsets in patient tumors. Boxes, median and interquartile range (IQR) of the ratios; whiskers, 1.5 IQR. **(F)** Signature of immune activation (SIA), defined as the ratio of CD8^+^ cell density to the sum of CD8^+^ and M2-like cell densities.

First, we evaluated the prognostic impact of the densities of the different immune cells in therapy naïve stage I-III colon cancers (*n*=286). Two cell classes demonstrated association of cell density with overall survival (OS), namely CD8^+^ T lymphocytes (positive association, *p*=0.042) and M2-like macrophages (negative association, *p*=0.004) (Fig 1B). Neither the pan-macrophage marker CD68 alone nor CD163, which is considered a marker of M2 differentiation, alone were associated with survival, whereas a more stringently defined M2-like macrophage subset expressing both CD68 and CD163 was (Fig 1B and C). Across all tumors, the CD68^+^CD163^+^ M2-like macrophages constituted only 5% of the CD68^+^ macrophages and 23% of the CD163^+^ cells (Fig 1D), but demonstrated substantial inter-patient heterogeneity with cell density ranging from 0 to 1080 cells/mm^2^ of tumor tissue (Fig 1E). Hypothesizing that these two immune cell types capture the interplay between anti- and pro-tumoral aspects of the immune microenvironment, we created a signature of immune activation (SIA) based on the relative infiltration levels of CD8^+^ cells and M2-like macrophages (Fig 1F and Supplementary materials).

To determine the prognostic value of SIA with regard to OS and recurrence-free survival (RFS), we transformed into a three-level categorized variable, using an unbiased approach with 33.3 and 66.6 percentiles as cutoffs. For comparison, we generated an Immunoscore-like metric (IS) by quantifying densities of CD3^+^ and CD8^+^ cells at the tumor center and invasive margin (*4*). Both IS and SIA demonstrated strong associations with OS and RFS in colon cancer stage I-III (Fig 2A and B). Interestingly, in a multivariate Cox model adjusted for pT stage, pN stage, patient age, gender and MSI status, both SIA and IS were independent predictors for OS and RFS (Table 1). Next, we compared the predictive ability of SIA to IS and well-known clinical risk factors. Integrative time-dependent estimation of the area under receiver-operator curve (iAUC) identified T stage as the strongest clinical predictor for OS (median iAUC 0.58) and N stage for RFS (median iAUC 0.58) (Fig 2C). However, both clinical risk factors and IS were inferior to SIA (median iAUC 0.59 for OS and RFS). Adding SIA to the model with combined clinical parameters improved the predictive ability (median iAUC 0.66 and 0.67 for OS and RFS). Finally, integration of clinical parameters, IS and SIA in one model resulted in median iAUC 0.68 and 0.69 for OS and RFS, respectively. The relative contribution to OS prediction was higher for SIA than for T and N stage, and when including IS in the model, the relative contribution of SIA and IS exceeded 50% and clearly surpassed the known clinical factors (Fig 2D). Because of the clinical need to identify high-risk tumors in stage II colon cancer patients, we analyzed this patient subgroup separately (*n*=117) and observed similar results with SIA stratifying high and low-risk disease (Fig S1A). Next, we assessed CRC patients with metastatic disease (*n*=66), stratified into three equally-sized terciles according to SIA and observed longer OS in the SIA-high group (Fig S1B). Thus, SIA demonstrated independent prognostic performance superior to the strongest clinical predictors (T and N stage), added substantial value to the multivariate prediction model in colon cancer patients of stages I-III, and demonstrated prognostic ability in stage II colon cancer and in metastatic CRC patients.

**Fig. 2.**
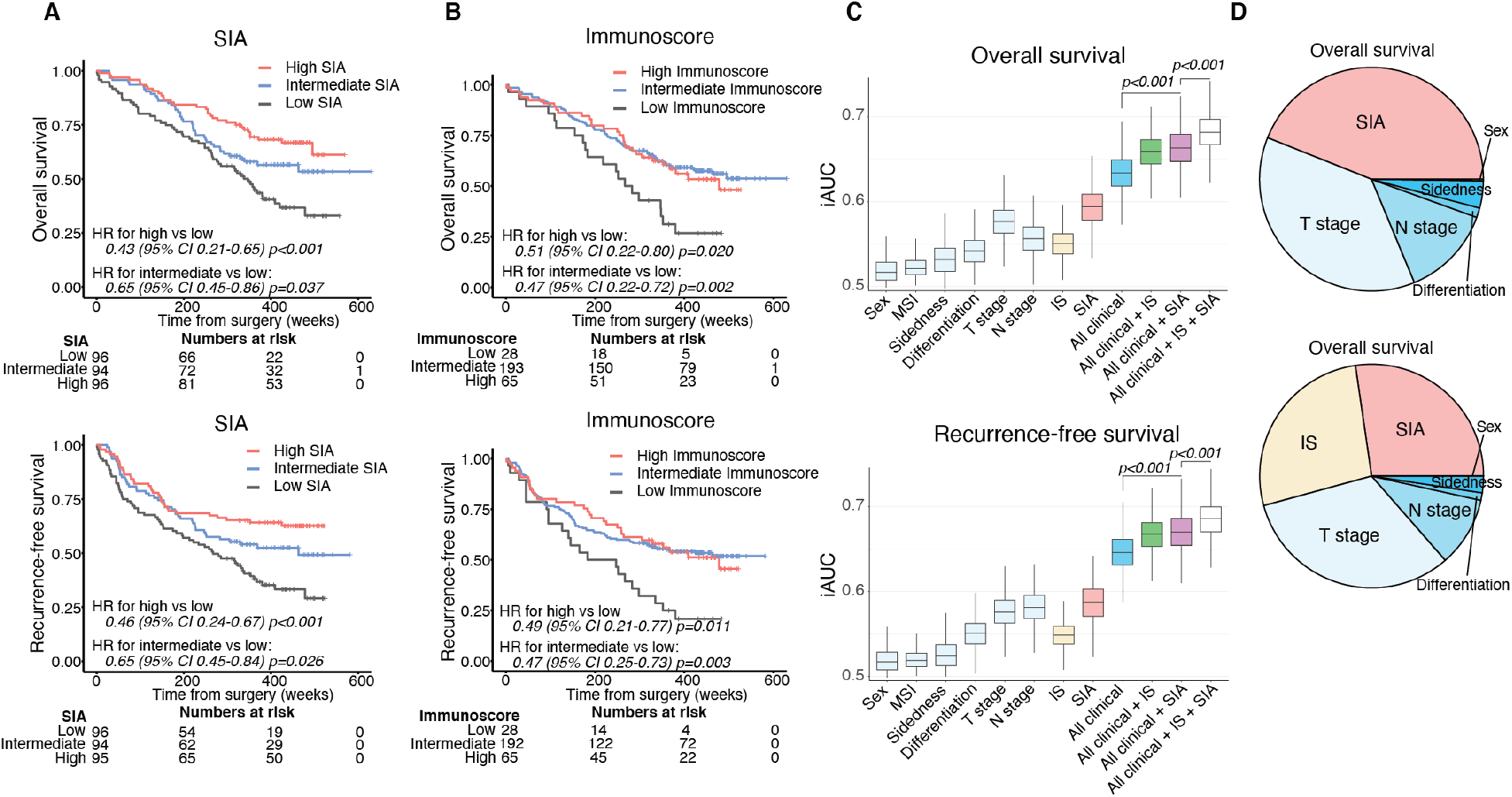
The SIA is an independent prognostic marker with performance superior to established clinical and immunological predictors for overall (OS) and recurrence-free (RFS) survival in therapy-naïve colon cancer stage I-III patients. **(A)** OS (upper panel) and RFS (lower panel) for the patients (n=286), stratified into SIA-low, -intermediate and -high groups, with SIA-low used as reference group. **(B)** OS (upper panel) and RFS (lower panel) for the patients stratified by trichotomized IS. Relative hazards were estimated by Cox proportional hazards model in (A) and (B). **(C)** Predictive accuracy of SIA, IS and clinical parameters for OS (upper panel) and RFS (lower panel) using iAUC analysis with 1000-fold bootstrap resampling. Univariate Cox proportional hazards models were applied to each of the analyzed factors separately and multivariate models used to evacuate the impact of factor combinations. **(D)** Relative contribution to the prediction of OS of SIA and clinical parameters (upper) or SIA, IS and clinical parameters (lower) determined using the χ^2^ proportion test.

**Table 1.**
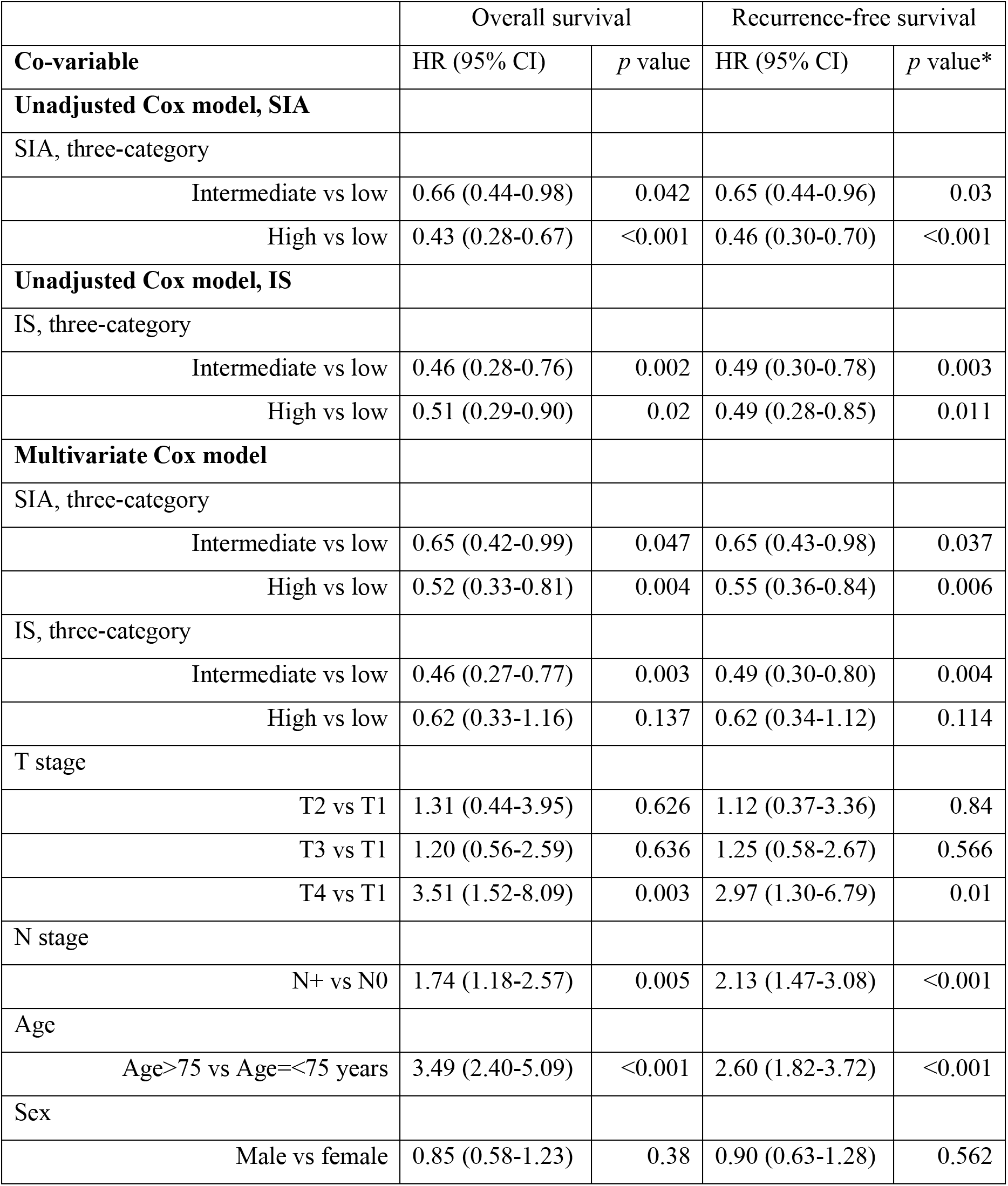
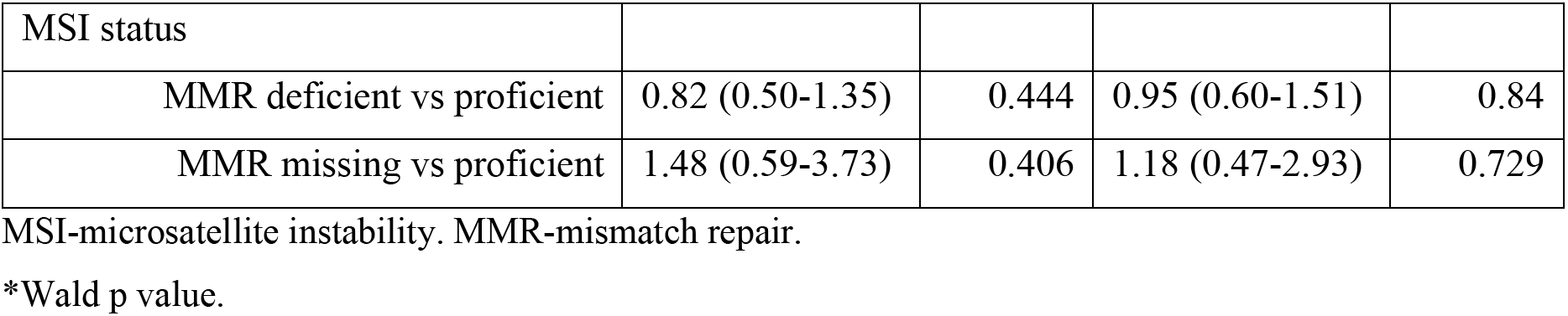
Strong prognostic capacity of SIA in predicting OS and RFS in therapy-naïve stage I-III colon cancer patients. Relative hazards, estimated in univariate (for SIA and IS separately) and multivariate (SIA, IS and clinical risk factors) Cox proportional hazards models, using OS and RFS as the endpoints.

Finally, we asked whether SIA was prognostic also in other cancers. We hypothesized that SIA would likely be of highest utility in tumor types with strong immunogenic properties (*13*). We ranked tumor types according to the number of mutations and neoantigens (Fig 3A), and analyzed four cohorts of tumors characterized by high counts, namely melanoma (*14*), lung carcinoma (*15*), bladder urothelial cancer (*16, 17*) and gastroesophageal adenocarcinomas (*18–20*). We also included two tumor types with low mutation and neoantigen density, endometrial (*21, 22*) and ovarian cancer (*23, 24*). Patients were stratified in terciles according to SIA, except for melanoma where the median was used since 41% of patients had the highest possible SIA value. High SIA was associated with longer survival in the four tumor types with high mutation and neoantigen count (*p*=0.001-0.037) while no association was seen in endometrial (*p*=0.996) and ovarian (*p*=0.399) cancers (Fig 3B). Further, SIA surpassed IS for prediction of OS in the four cohorts, demonstrating median iAUC ranging from 0.55 in bladder cancer to 0.61 in melanoma (Fig 3C). Thus, the SIA is a prognostic factor in at least five tumor types.

**Fig. 3.**
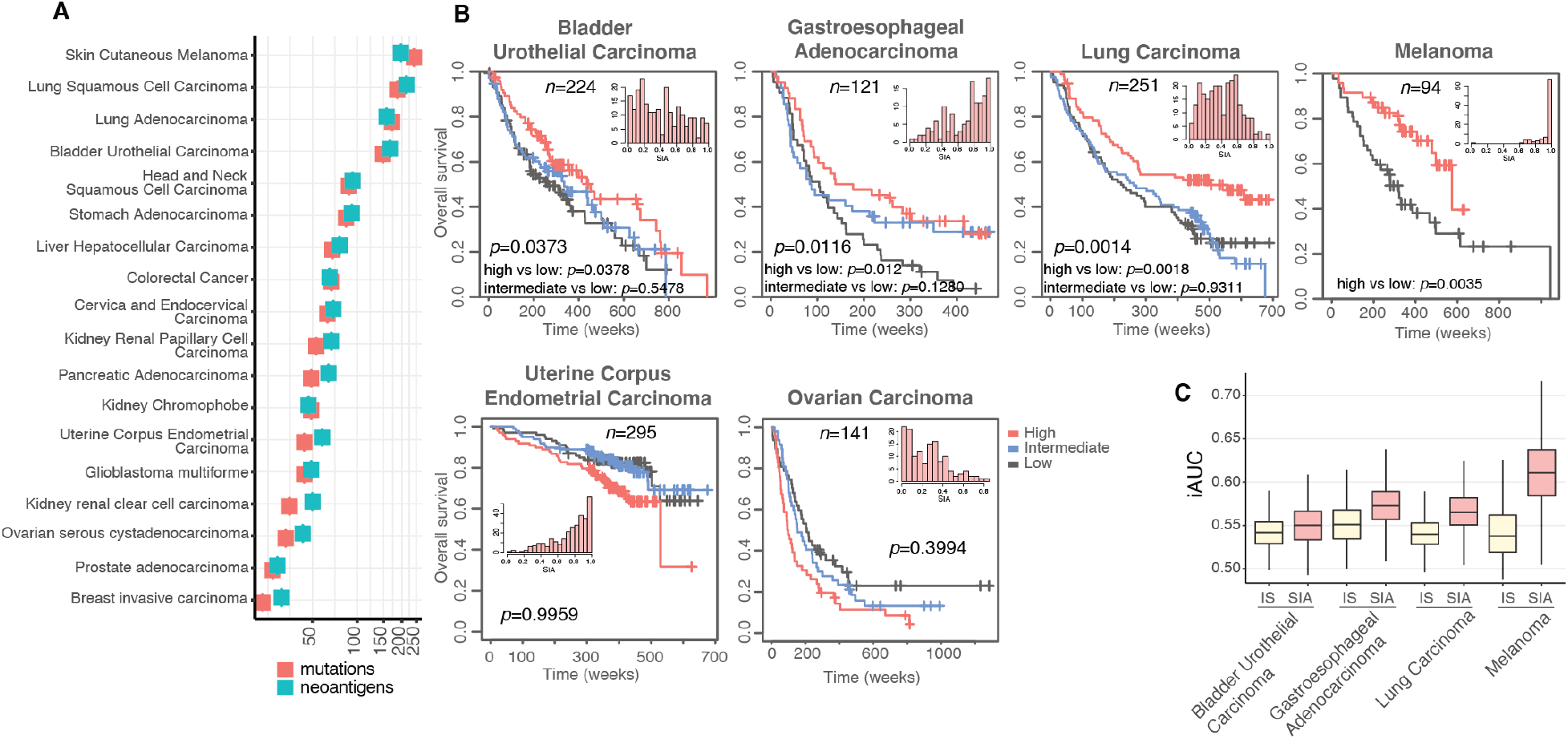
The SIA is prognostic in bladder cancer, cancer of the gastroesophageal junction, lung cancer and melanoma. **(A)** Median numbers of mutations and neoantigens across 19 solid cancers. The data was obtained from The Cancer Immunome Atlas (TCIA) (*27*). **(B)** Overall survival stratified by SIA in six tumor types. Tissue microarrays encompassing 91-295 cases of the respective tumor type were stained and the patients in each cohort stratified in terciles according to SIA score, except melanoma, which was stratified in two groups split by the median. Statistical analysis by Cox regression. Insets, histograms of density plots demonstrating the distribution of the continuous SIA metric. **(C)** Predictive accuracy of IS and SIA for OS in four cancer cohorts, generated using iAUC analysis with 1000-fold bootstrap resampling.

In summary, by immune cell sub-classification we confirmed the prognostic impact of CD8^+^ cell infiltration and revealed a prognostic subset of macrophages that was undetectable using a single-marker approach. The relationship of CD68^+^CD163^+^ macrophages polarized towards the M2 phenotype to anti-tumoral CD8^+^ cells provided a prognostic metric for the balance between pro- and anti-tumoral immunity. As both SIA and the Immunoscore-like score were independent variables in the multivariate analysis, these two metrics presumably reflect different aspects of tumor immunity. The SIA, unlike Immunoscore, does not require independent assessment of the tumor central region and invasive margin, and is prognostic in at least five tumor types. Given the ~6.7×10^6^ new cases and >4.3×10^6^ cancer-related deaths annually in these five cancers (*25*), the SIA has potential to enhance the reliability of prognosis prediction, improve the identification of high-risk patients and improve therapy decisions for >4×10^6^ patients per year, thus motivating the introduction of a TNM-Immune cancer classification (*26*).

## Acknowledgments

The collection of the colorectal cancer material and TMA production was supported by U-CAN, through Uppsala Biobank and the Department of Clinical Pathology, Uppsala University Hospital.

## Author contributions

Conceived study: A.M., T.S.; Designed experiments: A.M., K.L. B.G., P.M., T.S.; Performed experiments: A.M., M.B., S.S.; Image curation: I.H., S.M., S.K.; TMA-cohort construction: P.-U.M. (bladder); J.B. (lung), B.N., K.J., F.P. (CRC, melanoma), P.M. (lung) Patient database curation: J.M., K.H., M.A. (melanoma), P.-H.E., J.H. (endometrial), U.S. (bladder), P.-U.M. (bladder), J.B. (lung), C.H. (gastroesophageal), D.B. (gastroesophageal), J.B. (ovarian), H.S. (ovarian), K.J., F.P.; Data analysis: A.M., A. M.-B., J.E.; Data interpretation: A.M., P.M., T.S.; Wrote the manuscript: A.M., C.L, K.L., B.G., A.P., P.M., T.S.

## Data and materials availability

Data regarding methodology, image analysis, curation and data processing, and raw-data of stroma fraction is available from the corresponding author.

## Supplementary Materials

Materials and methods

Figure S1

Tables S1-S3

References (*4, 11, 12, 14-16, 18-24, 27*)

