## Supplementary file for "The ratio of cytotoxic lymphocytes to M2-like macrophages is prognostic in immunogenic tumors"

**This PDF file includes:**

Materials and Methods
Fig. S1
Tables S1 to S3

### Materials and Methods

#### Study cohorts and tissue microarrays

*The colorectal cancer (CRC) cohort* consists of prospectively collected CRC patients living in Uppsala County, Sweden, most of whom have been included in the Uppsala-Umeå Comprehensive Cancer Consortium (U-CAN) (*27*). In total, 937 patients were diagnosed with CRC between 2010 and 2014 in the Uppsala region. Of them, 746 (80%) were included in a TMA. For the present study, only patients with TMA material from primary tumors were selected. After the staining procedures and quality control, 497 patients had data from both immune panels (see below in *Multiplex immunofluorescence staining* for more details) of whom 286 patients had TNM stage I-III therapy naïve colon cancer. The clinicopathological characteristics of the included patients and their tumors are presented in Table S1. All patients received stage-stratified standard of care according to the Swedish national guidelines from 2008. According to the guidelines, colon tumors were recommended primary surgery and adjuvant chemotherapy if risk factors for recurrence were present. If the colon tumor was considered inextirpable/borderline resectable, neoadjuvant chemotherapy was administered to shrink the tumor before surgery. Rectal cancers received preoperative or neo-adjuvant radiotherapy/chemoradiotherapy stratified according to risk for locoregional or systemic recurrence. Formalin-fixed paraffin-embedded tissue blocks of primary tumors and distant metastases were used to construct TMAs. Each case was represented on the TMA with cores derived from the central part of the tumor and from the invasive margin. The study was performed within ethical permits from the regional ethical committee in Uppsala, Sweden (2010/198 and 2015/419).

The *melanoma* cohort encompassed TMA cores from 94 patients diagnosed with primary cutaneous malignant melanoma in the Uppsala region, Sweden, from 1980 to 2004 (*14*) (Table S2). The study was approved by the regional ethical committee in Uppsala, Sweden (2005/232).

The *lung cancer* cohort encompassed TMA cores from 251 patients diagnosed with Non-Small Cell Lung Cancer who underwent surgical treatment at Uppsala University Hospital, Sweden from 2006 to 2010 (*15*) (Table S2). The study was performed under a permit from the regional ethical committee in Uppsala (2012/532).

The *gastroesophageal cancer* cohort included TMA cores from 121 patients with chemoradiotherapy-naïve gastroesophageal adenocarcinomas who underwent surgery at the University Hospitals of Lund and Malmö from 2006 to 2010 (*18-20*) (Table S2). The study was performed under a permit from the regional ethical committee in Lund (2007/445).

The *urothelial cancer* cohort encompassed TMA cores collected from primary urothelial tumors from 224 patients undergoing surgery at Uppsala University Hospital between 1984 and 2005 (*16*) (Table S2). The study was performed under a permit from the regional ethical committee in Uppsala (2005/143).

The *uterine corpus endometrial carcinoma* cohort consisted of TMA cores from 295 uterine carcinomas from patients surgically treated at Turku University Hospital, Finland, between 2004-2007 (*21, 22*) (Table S2). The study was performed under a permit from the ethical review board in Helsinki (2016/010).

The *ovarian carcinoma* cohort was presented as TMA cores from invasive ovarian cancer cases, derived from two pooled prospective, population-based cohorts; the Malmö Diet and Cancer Study and the Malmö Preventive Project (*23, 24*) (Table S2). The study was performed under a permit from the regional ethical committee in Lund (2007/445).

**Multiplex immunofluorescence staining**

For the multiplexed immunofluorescence staining, 4 μm thick TMA sections were de-paraffinized, rehydrated and rinsed in distilled $H_2O$. Two staining protocols were established for the two panels of antibodies: the lymphocyte panel, with CD4, CD8, CD20, FoxP3, CD45RO, and pan-cytokeratin (CK), and the NK/macrophage panel encompassing CD56, NKp46, CD3, CD68, CD163, and pan-CK. The staining procedure was performed as described (*11, 12*). Detailed staining conditions and reagent references are provided in Table S3.

**Imaging, image analysis, and thresholding**

The stained TMAs were imaged using the Vectra Polaris system (Akoya) in multispectral mode at a resolution of 2 pixels/μm. Each of the images was manually reviewed and curated by a pathologists with a support from HistoOne AB (Uppsala, Sweden), to exclude artefacts, staining defects and accumulation of immune cells in necrotic areas and intraglandular structures. The vendor-provided machine learning algorithm, implemented in the inForm software, was trained to split tissue into three categories: tumor compartment, stromal compartment, or blank areas. The training was performed for each cohort separately by providing a set of the samples that was manually annotated by pathologists. Cell segmentation was performed using DAPI nuclear staining as described (*11, 12*). The perinuclear region at 3 μm (6 pixels) from the nuclear border was considered the cytoplasm area. The cell phenotyping function of the inForm software was used to manually define a representative subset of cells positive to expression of each of the markers and a subset of cells negative to all markers. The intensity of the marker expression in selected cells was used to set the thresholds for marker positivity.

Intensity thresholds for the markers were determined in the R programming environment [R Core Team, 2013] by GeneVia Technologies (Tampere, Finland). The marker-specific thresholds were defined by the distributions of the positive and negative cell intensities for that marker. Marker-specific probability density distributions were estimated by smoothing the intensity values with Gaussian kernel estimation with automatic bandwidth detection using the density function of the R package *stats*. The intensity thresholds for each marker were established as (1) the mean value of the highest intensity of the negative cells and the lowest intensity of the positive cells, if the intensities of the positive and negative cells did not overlap, or (2) as the intensity value which minimized the overall classification error based on the probability density distributions, if there was overlap. The False Positive Rate, True Positive Rate, False Negative Rate, True Negative Rate, and the overall classification error were calculated for each established threshold, i.e. for each marker, and controlled individually. The thresholds were established separately and independently for each tumor type and were applied to the raw output data of the complete cohorts. Every cell was thus characterized as positive or negative for each marker in the panel. This data was used to classify the cell and define its immune subtype (Fig 1A). Finally, cell counts were normalized against analyzed tissue area size and used as cell density (units per $mm^2$) in further analyses.

**Signature of immune activation and Immunoscore**

The signature of immune activation (SIA) was computed as a ratio of CD8+ cell density to the sum of the densities of CD8+ and M2-like cells, or SIA=(CD8 density)/(CD8 density+M2-like density). For the Immunoscore (IS), each tumor in the CRC TMA cohort was represented by TMA cores derived from the central part and the invasive margin of the tumors. The CD3 and CD8-positive cells were defined in each of the regions, thus resulting in four values per case (i.e. CD3 density in tumor center, CD8 density in tumor center, CD3 density at the invasive margin, CD8 density at the invasive margin). The IS was generated as described (*4*) by computing a mean of the four. In the other cohorts, the TMA cores were obtained from the bulk tumor region, without separation between central parts and invasive margin. Thus, for these tumors two values per case were obtained (CD3 and CD8-positive cell density) and IS was generated by computing a mean of the two. Further, using the mean percentiles, IS was categorized into 3 groups: Low (mean percentile 0-25%), Intermediate (25-70%) and High (70-100%).

**Mutations and Neoantigens**

The median values of the numbers of mutations and neoantigens across 19 solid cancers were obtained from The Cancer Immunome Atlas (TCIA) project (https://tcia.at/home).

**Statistics**

Statistical analyses were performed using R (version 3.5.1) and SPSS V20 (SPSS Inc., Chicago, IL). In radically operated CRC stage I-III patients, recurrence-free survival (RFS) was computed as the time from surgery to the first documented disease progression including local recurrence or distant metastases or death due to any reason, whichever occurred first. Overall survival (OS) was the time from surgery to death due to any reason. To estimate relative hazards in both univariate and multivariable models, a Cox proportional hazards model was used. The predictive accuracies of the models were evaluated by 1000-fold bootstrap resampling and by computing the time-dependent area under the receiver operating characteristic curve (iAUC) for each bootstrap sampling. The relative importance of parameters for the estimation of survival risk was computed using the chi squared proportion ($\chi^2$).

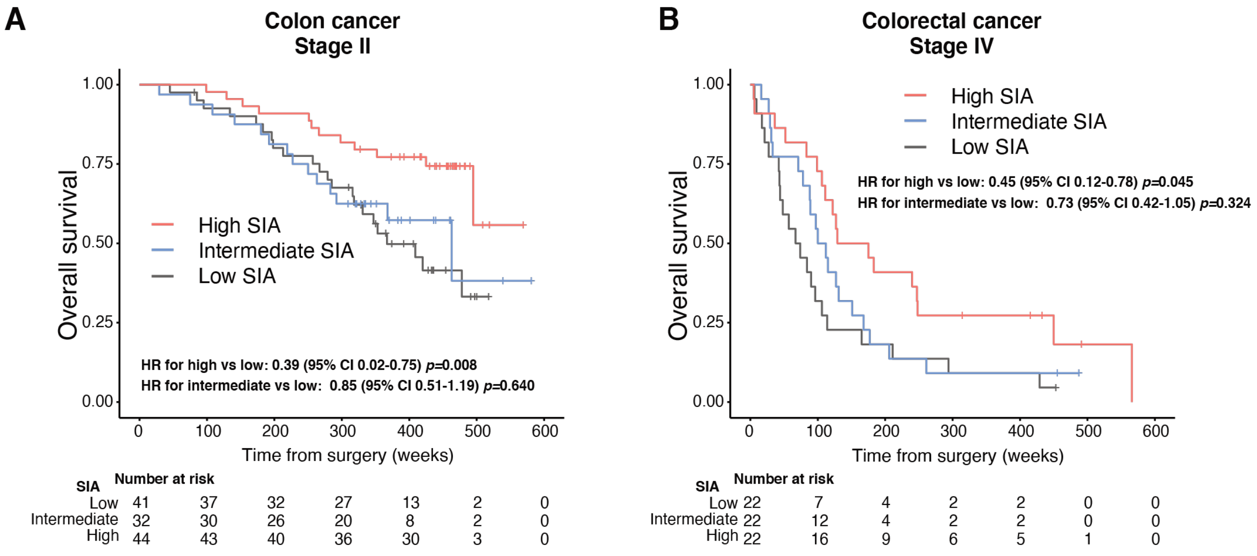


**Fig. S1. The SIA is prognostic in (A) therapy-naïve colon cancer stage II patients and (B) metastatic colorectal cancer patients**. Kaplan-Meier curves and numbers at risk demonstrate OS in patient groups, stratified by trichotomized SIA. Cox proportional hazards models were used to estimate relative hazards.

**Table S1. Baseline clinicopathological characteristics of the colorectal cancer cohort.** Patient data shown for cases where successful staining was available from the TIL panel, the NK/MF panel or where data from both panels were available and SIA was calculated. Values are shown as the number of cases (percentages) unless indicated otherwise. Percentages may not add to 100% due to rounding. MSI, microsatellite instability; MMR, mismatch repair; RT, radiotherapy; scRT, short-course radiotherapy; lcRT, long-course radiotherapy; CT, chemotherapy; CRT, chemo-radio therapy.

| **Characteristics** | TIL Panel (n=520) | NK/MF Panel (n=508) | SIA group (n=497) |
|---|---|---|---|
| **Age** | | | |
| Average age in years ± SD | 69.83 ± 12.02 | 69.76 ± 11.89 | 69.86 ± 11.86 |
| < 75 and = 75 years old | 338 (65%) | 331 (65.2%) | 323 (65%) |
| > 75 years old | 182 (35%) | 117 (34.8%) | 174 (35%) |
| **Sex** | | | |
| Male | 282 (54.2%) | 274 (53.9%) | 268 (53.9%) |
| Female | 238 (45.8%) | 234 (46.1%) | 229 (46.1%) |
| **Location** | | | |
| Colon | 365 (70.2%) | 355 (69.9%) | 351 (70.6%) |
| Rectum | 155 (29.8%) | 153 (30.1%) | 146 (29.4%) |
| **pT Stage** | | | |
| 0 | 3 (0.6%) | 5 (1%) | 2 (4%) |
| 1 | 39 (7.5%) | 37 (7.3%) | 37 (7.4%) |
| 2 | 61 (11.7%) | 60 (11.8%) | 58 (11.7%) |
| 3 | 309 (59.4%) | 304 (59.8%) | 300 (60.4%) |
| 4 | 99 (19%) | 95 (18.7%) | 93 (18.7%) |
| Missing data | 9 (1.7%) | 7 (1.4%) | 7 (1.4%) |
| **pN Stage** | | | |
| 0 | 273 (52.5%) | 266 (52.4%) | 261 (52.5%) |
| 1 | 237 (45.6%) | 236 (46.5%) | 230 (46.3%) |
| Missing data | 10 (1.9%) | 6 (1.2%) | 6 (1.2%) |
| **pM Stage** | | | |
| 0 | 437 (84%) | 426 (83.9%) | 418 (84.1%) |
| 1 | 77 (14.8%) | 77 (15.2%) | 74 (14.9%) |
| Missing data | 6 (1.2%) | 5 (1%) | 5 (1%) |
| **pTNM stage** | | | |
| 0 | 4 (0.8%) | 5 (1%) | 3 (0.6%) |
| 1 | 85 (16.3%) | 82 (16.1%) | 80 (16.1%) |
| 2 | 172 (33.1%) | 166 (34.6%) | 165 (33.2%) |
| 3 | 179 (34.4%) | 176 (34.6%) | 173 (34.8%) |
| 4 | 77 (14.8%) | 77 (15.2%) | 74 (99.6%) |

| | | | |
|---|---|---|---|
| Missing data | 3 (0.6%) | 2 (0.4%) | 2 (0.4%) |
| **Differentiation Grade** | | | |
| Low | 368 (70.8%) | 356 (70.1%) | 353 (71%) |
| High | 84 (16.2%) | 83 (16.3%) | 81 (16.3%) |
| Missing data | 68 (13.1%) | 69 (13.6%) | 63 (12.7%) |
| **Neural Invasion** | | | |
| No | 361 (69.4%) | 352 (69.3%) | 343 (69%) |
| Yes | 80 (15.4%) | 78 (15.4%) | 78 (15.7%) |
| Missing data | 79 (15.2%) | 78 (15.4%) | 76 (15.3%) |
| **Vascular Invasion** | | | |
| No | 132 (25.4%) | 326 (64.2%) | 317 (63.8%) |
| Yes | 331 (63.7%) | 125 (24.6%) | 125 (25.2%) |
| Missing data | 57 (11%) | 57 (11.2%) | 55 (11.1%) |
| **MSI/MMR Status** | | | |
| MMR proficient | 441 (84.8%) | 432 (85%) | 424 (85.3%) |
| MMR deficient | 66 (12.7%) | 64 (12.6%) | 62 (12.5%) |
| Missing data | 13 (2.5%) | 12 (2.4%) | 11 (2.2%) |
| **BRAF Mutation** | | | |
| No | 206 (39.6%) | 201 (39.6%) | 197 (39.6%) |
| Yes | 46 (8.8%) | 45 (8.9%) | 45 (9.1%) |
| Missing data | 268 (51.5%) | 262 (51.5%) | 255 (51.3%) |
| **Neoadjuvant Treatment (in Rectal only)** | | | |
| No | 51 (32.9%) | 48 (31.4%) | 47 (32.3%) |
| Yes | 91 (58.7%) | 93 (60.8%) | 87 (59.6%) |
| Missing data | 13 (8.4%) | 12 (7.8%) | 12 (8.2%) |
| **Neoadjuvant Treatment Type (in Rectal only)** | | | |
| scRT+delay | 32 (20.6%) | 31 (20.3%) | 31 (21.2%) |
| scRT+immediate | 29 (18.7%) | 28 (18.3%) | 28 (19.2%) |
| scRT+CT | 10 (6.5%) | 11 (7.2%) | 9 (6.2%) |
| CRT | 8 (5.2%) | 10 (6.5%) | 8 (5.5%) |
| CT | 1 (0.6%) | 1 (0.7%) | 0 |
| lcRT | 1 (0.6%) | 1 (0.7%) | 1 (0.7%) |
| Missing data | 22 (14.2%) | 22 (14.2%) | 21 (14.4%) |

**Table S2. Baseline clinicopathological characteristics of validation cohorts.** Data from cases where SIA could be computed. Values are shown as the number of cases (percentages) unless indicated otherwise. Percentages rounded to one decimal. *Median survival times were calculated using the Kaplan-Meier method. Mean survival times were estimated when median survival times cannot be calculated from the data

| | Endometrial Cancer | Non Small Cell Lung Cancer | Urine Bladder Cancer | Ovarian Cancer | Gastro-esophageal Cancer | Melanoma |
|---|---|---|---|---|---|---|
| **Patient sample size and median survival time** | | | | | | |
| Total number of patients (%) | 295 (26.1%) | 251 (22.2%) | 224 (19.8%) | 141 (12.5%) | 127 (11.2%) | 94 (8.3%) |
| Median survival time* ± SD (weeks) | 545.08 ± 14.19[b] | 276 ± 43.58 | 348 ± 50.85 | 151 ± 18.55 | 113 ± 20.80 | 478.4 ± 56.96 |
| **Age at diagnosis** | | | | | | |
| Mean ± SD | 66 ± 10.47 | 67.19 ± 7.45 | 70.69 ± 11.81 | 63.70 ± 8.39 | 70.8 ± 11.04 | 60.45 ± 14.61 |
| Median ± SD | 66 ± 10.47 | 67 ± 7.45 | 72 ± 11.81 | 63 ± 8.39 | 72 ± 11.04 | 61.09 ± 14.61 |
| ≤ Median | 152 (51.5%) | 132 (52.6%) | 115 (51.3%) | 73 (51.8%) | 64 (50.4%) | 47 (50%) |
| > Median | 143 (48.5%) | 119 (47.4%) | 109 (48.7%) | 68 (48.2%) | 63 (49.6%) | 47 (50%) |
| **Sex** | | | | | | |
| Female | 295 (100%) | 128 (51%) | 50 (22.3%) | 141 (100%) | 29 (22.8%) | 44 (46.8%) |
| Male | - | 123 (49%) | 174 (77.7%) | - | 98 (77.2%) | 50 (53.2%) |
| **pT classification** | | | | | | |
| pT0 | 0 | 0 | 102 (45.5%) | 0 | 0 | 0 |
| pT1 | 0 | 113 (45%) | 95 (42.4%) | 0 | 7 (5.5%) | 23 (24.5%) |
| pT2 | 0 | 102 (40.6%) | 18 (8%) | 0 | 22 (17.3%) | 27 (28.7%) |
| pT3 | 0 | 28 (11.2%) | 7 (3.1%) | 0 | 77 (60.6%) | 23 (24.5%) |
| pT4 | 0 | 8 (3.2%) | 2 (0.9%) | 0 | 21 (16.5%) | 15 (16%) |
| Missing data | 295 (100%) | 0 | 0 | 141 (100%) | 0 | 6 (6.4%) |
| **pN classification** | | | | | | |
| pN0 | 0 | 193 (76.9%) | 24 (10.7%) | 0 | 36 (28.3%) | 0 |
| pN1 | 0 | 29 (11.6%) | 5 (2.2%) | 0 | 26 (20.5%) | 0 |
| pN2 | 0 | 29 (11.6%) | 0 | 0 | 33 (26%) | 0 |
| pN3 | 0 | 0 | 0 | 0 | 32 (25.2%) | 0 |
| Missing data | 295 (100%) | 0 | 195 (87.1%) | 141 (100%) | 0 | 94 (100%) |
| **pM classification** | | | | | | |
| pM0 | 0 | 0 | 60 (26.8%) | 0 | 116 (91.3%) | 0 |
| pM1 | 0 | 0 | 18 (8%) | 0 | 3 (2.4%) | 0 |
| pM2 | 0 | 0 | 0 | 0 | 8 (6.3%) | 0 |
| Missing data | 295 (100%) | 251 (100%) | 146 (65.2%) | 141 (100%) | 0 | 94 (100%) |

| | | | | | | |
|---|---|---|---|---|---|---|
| **Clinical stage at diagnosis** | | | | | | |
| 1 | 242 (82%) | 155 (61.8%) | 0 | 23 (16.3%) | 0 | 0 |
| 2 | 8 (2.7%) | 53 (21.1%) | 0 | 16 (11.3%) | 0 | 0 |
| 3 | 39 (13.2%) | 36 (14.3%) | 0 | 72 (51.1%) | 0 | 0 |
| 4 | 6 (2%) | 7 (2.8%) | 0 | 19 (13.5%) | 0 | 0 |
| Missing data | 0 | 0 | 224 (100%) | 11 (7.8%) | 127 (100%) | 94 (100%) |
| **Differentiation grade / Histological differentiation** | | | | | | |
| G1 Well differentiated (Low grade) | 247 (83.7%) | 0 | 71 (31.7%) | 4 (2.8%) | 77 (60.6%) | 0 |
| G2 Moderately differentiated (Intermediate grade) | - | - | - | 31 (22%) | - | - |
| G3 Poorly differentiated (High grade) | 48 (16.3%) | 0 | 153 (68.3%) | 106 (75.2%) | 50 (39.4%) | 0 |
| Missing data | 0 | 251 (100%) | 0 | 0 | 0 | 94 (100%) |
| **WHO performance status** | | | | | | |
| 0 | 0 | 153 (61%) | 0 | 0 | 0 | 0 |
| 1 | 0 | 95 (37.8%) | 0 | 0 | 0 | 0 |
| 2 | 0 | 3 (1.2%) | 0 | 0 | 0 | 0 |
| ≥ 3 | 0 | 0 | 0 | 0 | 0 | 0 |
| Missing data | 295 (100%) | 0 | 224 (100%) | 141 (100%) | 127 (100%) | 94 (100%) |
| **Resection margin** | | | | | | |
| R0 | 0 | 0 | 0 | 0 | 85 (66.9%) | 0 |
| R1 | 0 | 0 | 0 | 0 | 35 (27.6%) | 0 |
| R2 | 0 | 0 | 0 | 0 | 7 (5.5%) | 0 |
| Missing data | 295 (100%) | 251 (100%) | 224 (100%) | 141 (100%) | 0 | 94 (100%) |
| **p53 status** | | | | | | |
| Mutant | 32 (10.8%) | 0 | 0 | 0 | 0 | 0 |
| Wild-type | 263 (89.2%) | 0 | 0 | 0 | 0 | 0 |
| Missing data | 0 | 251 (100%) | 224 (100%) | 141 (100%) | 127 (100%) | 94 (100%) |
| **MSI/MSS status** | | | | | | |
| MSI | 0 | 0 | 0 | 2 (1.4%) | 11 (8.7%) | 0 |
| MSS | 0 | 0 | 0 | 136 (96.5%) | 116 (91.3%) | 0 |
| Missing data | 295 (100%) | 251 (100%) | 224 (100%) | 3 (2.1%) | 0 | 94 (100%) |
| **Neural invasion** | | | | | | |
| Yes | 0 | 0 | 0 | 0 | 9 (7.1%) | 0 |
| No | 0 | 0 | 0 | 0 | 22 (17.3%) | 0 |
| Missing data | 295 (100%) | 251 (100%) | 224 (100%) | 141 (100%) | 96 (75.6%) | 94 (100%) |
| **Vascular invasion** | | | | | | |
| Yes | 0 | 0 | 0 | 0 | 10 (7.9%) | 0 |

| | | | | | | |
|---|---|---|---|---|---|---|
| No | 0 | 0 | 0 | 0 | 41 (32.3%) | 0 |
| Missing data | 295 (100%) | 251 (100%) | 224 (100%) | 141 (100%) | 76 (59.8%) | 94 (100%) |
| **Smoking history** | | | | | | |
| Current smoker | 0 | 131 (52.2%) | 92 (41.1%) | 0 | 0 | 0 |
| Non-current smoker | 0 | 120 (47.8%) | 18 (8%) | 0 | 0 | 0 |
| Missing data | 295 (100%) | 0 | 114 (50.9%) | 141 (100%) | 127 (100%) | 94 (100%) |
| **Adjuvant treatment** | | | | | | |
| Yes | 0 | 0 | 2 (0.9%) | 0 | 0 | 0 |
| No | 0 | 251 (100%) | 0 | 0 | 0 | 0 |
| Missing data | 295 (100%) | 0 | 222 (99.1%) | 141 (100%) | 127 (100%) | 94 (100%) |
| **Neoadjuvant treatment** | | | | | | |
| Yes | 270 (91.5%) | 102 (40.6%) | 69 (30.8%) | 70 (49.6%) | 10 (7.9%) | 0 |
| No | 25 (8.5%) | 129 (51.4%) | 0 | 1 (0.7%) | 117 (92.1%) | 0 |
| Missing data | 0 | 20 (8%) | 115 (69.2%) | 70 (49.6%) | 0 | 94 (100%) |

**Table S3. Antibodies and amplification reagents used for multiplex fluorescent IHC.**

Staining protocols were established for the lymphocyte and the NK/macrophage panel, respectively. The staining procedure included 5 to 6 cycles of microwave treatment, incubation with primary antibody, and amplification system fluorophore labeling. The final cycle was followed by DAPI staining and slide mounting. *Antigen retrieval performed in microwave oven at 100 °C, 15min. †Amplification systems ImmPRESS® HRP or Opal HRP were used: The ImmPRESS® HRP Anti-Mouse IgG (Peroxidase) (Cat. No: MP-7402-50) and Anti-Rabbit IgG (Peroxidase) Polymer Detection Kits, made in Horse (Cat No: MP-7401-50) (Vector Laboratories); Opal™ Polymer anti-Rabbit+anti-Mouse HRP Kit (Cat No: ARH1001EA) (Akoya). [#]in melanoma instead of cytokeratin/E-cadherin cocktail, Melan A was used to identify malignant tissue vs stroma.

| Panel | Order | Antigen retrieval* | Marker | Clone | Host Species | Dilution | Amplification/enzyme reagent† | Company |
|---|---|---|---|---|---|---|---|---|
| Lymphocyte panel | 1. | pH9 | CD8a | C8/144B | Mouse | 1:250 | ImPress | Thermo Fisher |
| | 2. | pH9 | CD4 | 4B12 | Mouse | 1:100 | ImPress | Agilent |
| | 3. | pH6 | CD20 | L26 | Mouse | 1:1500 | Opal HRP | Agilent |
| | 4. | pH6 | FoxP3 | D6O8R | Rabbit | 1:300 | ImPress | Cell Signaling |
| | 5. | pH6 | CD45RO | UCHL1 | Mouse | 1:400 | | Thermo Fisher |
| | 6. | pH6 | PanCK | C-11 | Mouse | 1:100 | Opal HRP | Abcam |
| | | | Cytokeratin | AE1/AE3 | Mouse | 1:400 | | Agilent |
| | | | E-cadherin | 36/E | Mouse | 1:2000 | | BD Biosciences |
| | 6.[#] | pH6 | Melan A | | | 1:100 | ImPress | |
| | 7. | - | DAPI | - | - | - | - | Akoya |
| NK/Macrophage panel | 1. | pH6 | CD3 | F7.2.38 | Mouse | 1:80 | ImPress | Agilent |
| | 2. | pH6 | CD56 | 123C3 | Mouse | 1:100 | ImPress | Agilent |
| | 3. | pH6 | NKp46 | NCR1 | Rabbit | 1:150 | ImPress | Thermo Fisher |
| | 4. | pH6 | CD68 | PG-M1 | Mouse | 1:100 | Opal HRP | Agilent |
| | 5. | pH6 | CD163 | 10D6 | Mouse | 1:400 | Opal HRP | Novocastra |
| | 6. | pH6 | PanCK | C-11 | Mouse | 1:100 | Opal HRP | Abcam |
| | | | Cytokeratin | AE1/AE3 | Mouse | 1:500 | | Agilent |
| | | | E-cadherin | 36/E | Mouse | 1:2000 | | BD Biosciences |
| | 6.[#] | pH6 | Melan A | | | 1:100 | ImPress | |
| | 7. | - | DAPI | - | - | - | - | Akoya |
